# A biologically accurate model of directional hearing in the parasitoid fly *Ormia ochracea*

**DOI:** 10.1101/2021.09.15.460520

**Authors:** Max R. Mikel-Stites, Mary K. Salcedo, John J. Socha, Paul E. Marek, Anne E. Staples

**Affiliations:** Department of Biomedical Engineering and Mechanics, Virginia Tech, Blacksburg, VA, 24061, USA; Engineering Mechanics program, Virginia Tech, Blacksburg, VA, 24061, USA; Department of Mathematics, Virginia Tech, Blacksburg, VA, 24061, USA; Department of Entomology, Virginia Tech, Blacksburg, VA, 24061, USA

**Keywords:** acoustics, binaural hearing, biomechanics, mathematical modeling, sensory modeling, *Ormia ochracea*, parasitoidism, bioinspiration

## Abstract

This manuscript was compiled on October 5, 2021 Although most binaural organisms localize sound sources using neurological structures to amplify the sounds they hear, some animals use mechanically coupled hearing organs instead. One of these animals, the parasitoid fly *Ormia ochracea,* has astoundingly accurate sound localization abilities and can locate objects in the azimuthal plane with a precision of 2°, equal to that of humans. This is accomplished despite an intertympanal distance of only 0.5 mm, which is less than 1/100th of the wavelength of the sound emitted by the crickets that it parasitizes. In 1995, Miles *et al.* developed a model of hearing mechanics in O. *ochracea,* which works well for incoming sound angles of less than ±30°, but suffers from reduced accuracy (up to 60% error) at higher angles. Even with this limitation, it has served as the basis for multiple bio-inspired microphone designs for decades. Here, we present critical improvements to the classic *O. ochracea* hearing model based on information from 3D reconstructions of *O. ochracea’s* tympana. The 3D images reveal that the tympanal organ has curved lateral faces in addition to the flat front-facing prosternal membranes represented in the Miles model. To mimic these faces, we incorporated spatially-varying spring and damper coefficients that respond asymmetrically to incident sound waves, making a new quasi-two-dimensional (q2D) model. The q2D model has high accuracy (average errors of less than 10%) for the entire range of incoming sound angles. This improved biomechanical hearing model can inform the development of new technologies and may help to play a key role in developing improved hearing aids.

**Significance Statement:** The ability to identify the location of sound sources is critical to organismal survival and for technologies that minimize unwanted background noise, such as directional microphones for hearing aids. Because of its exceptional auditory system, the parasitoid fly *Ormia ochracea* has served as an important model for binaural hearing and a source of bioinspiration for building tiny directional microphones with outsized sound localization abilities. Here, we performed 3D imaging of the fly’s tympanal organs and used the morphological information to improve the current model for hearing in *O. ochracea.* This model greatly expands the range of biological accuracy from ±30° to all incoming sound angles, providing a new avenue for studies of binaural hearing and further inspiration for fly-inspired technologies.

---

The ability to localize sound allows animals to avoid predators and assists them in finding mates and capturing prey. Binaural organisms, those with two ears, locate sound-emitting objects by comparing the intensity and timing of incident sound waves arriving at their two hearing organs (Fig. 1A). Sound localization in binaural organisms is commonly described using two metrics: 1) interaural time delay (ITD), the difference in time it takes sound to reach the two hearing organs, and 2) interaurual amplitude difference (IAD), the difference in sound amplitude between the two organs (Fig. 1A) (1).

**Fig. 1.**
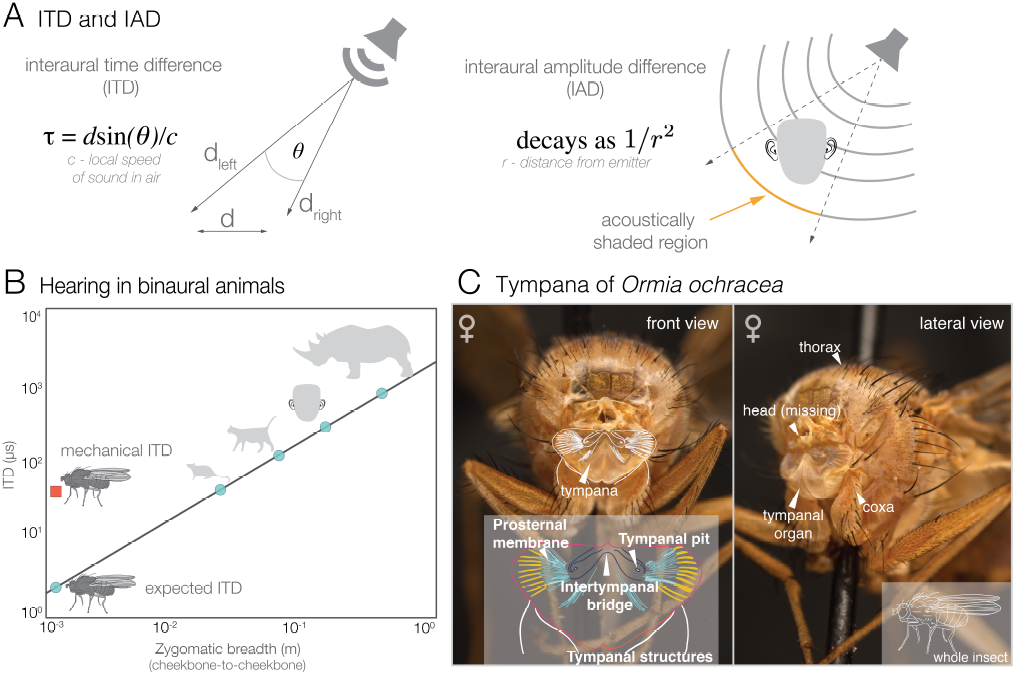
Hearing in binaural animals and the fly *Ormia ochracea*. **(A)** Physical meaning of ITD and IAD. ITD is the time delay between sound reaching one sensory organ relative to the other, defined by the equation *τ* = *dsin*(*θ*)/*c*, where *d* is the distance between the hearing organs and *c* is the speed of sound in air. IAD is the difference in response amplitude between the left and right sensory organs, due to acoustic shading or signal decay. **(B)** Approximate ITD values in representative animals (O. *ochracea,* rat, cat, human, and rhinoceros) for a sound source at a 45° angle from the midline, calculated using the formula in (A), with the zyogmatic breadth (cheekbone-to-cheekbone distance) used as an approximate measure of interaural distance for mammals. Data from (29–32). **(C)** Female *O. ochracea* post-decapitation, showing location of prosternal membrane, tympanal pits, and intertympanal bridge, key physical features in the modeling of its binaural hearing.

In vertebrates, ITD is calculated in the superior olivary nucleus of the brain stem and IAD is calculated in the inferior colliculus in the midbrain nucleus (2). By comparison, many invertebrates lack significant neural investment in central processing and rely heavily on mechanical structures to pre-process sensory signals (3–5). The particulars of sound localization are complex and vary widely among animals, as the ITD and IAD ranges experienced by binaural animals demonstrate (Fig. 1B) and the biophysics of sound localization for specific species are often too complex to be modeled well by simple analytical models. However, simplistic insect hearing models can be used to extract key principles of binaural hearing without complex physiological modeling of neural processes.

For the tachinid fly *Ormia ochracea,* the ability to hear its host plays a key role in its reproductive cycle and overall fitness. As a parasitoid, *O. ochracea* listens to chirping male crickets and follows the sound back to the source, where female *O. ochracea* then deposit their larvae (6). Gravid *O. ochracea* females will remain in an area for extended periods of time in response to cricket chirping sounds, even if no cricket is present (7). Given its small size, if *O. ochracea* relied exclusively on the distance between its prosternal tympanal membranes (Fig. 1C), the ITD it experienced would be at the nanosecond scale or below (far too small to reliably perceive differences in azimuthal angle between sound source locations), and there would be no practical difference in sound amplitude between the two membranes (IAD). To solve this scaling problem without resorting to bulky neurological investment, *O. ochracea* have two mechanically coupled membranous tympana directly beneath their head (Fig. 1C). These coupled tympana are composed of a pair of prosternal membranes, joined together by an intertympanal bridge (8), and are significantly larger in female *Ormia* (Fig. 1C). This distinctive mechanical coupling serves to increase the ITD and IAD perceived by the fly. We will refer to the increased ITD and IAD as mITD (mechanical ITD) and mIAD (mechanical IAD), respectively. These mechanically amplified values allow it to successfuly localize chirping crickets. Since *O. ochracea* are active at or after dusk, they also use their hearing to avoid predation by bats, exhibiting a startle response while in flight to bat sonar frequency sound, similar to preying mantises (9, 10). As such, the ability to accurately and quickly locate a source of incoming sound at high levels of lateral angular resolution is a significant advantage, especially in potentially noisy environments (11), suggesting that *O. ochracea* may be a good source of bioinspiration to tackle the so-called cocktail party problem (12) (isolating sounds in a noisy environment) for directional microphones and hearing aids.

Accurate models of binaural hearing in animals are generally highly complex. Because of the mechanical nature of its acoustic sensing organ, *O. ochracea* is one of the few exceptions, and it has been the focus of numerous studies featuring its uniquely “simple” hearing organs and how they function (8, 9, 13–19). To investigate the biomechanical mechanisms that underlie *O. ochracea*’s unusual hearing abilities, Miles, Robert, and Hoy developed mechanical and mathematical models of the ormiine’s coupled tympana in 1995 (15). The authors validated their model against experimental data, recording tympanal membrane positions and velocities, and consequently mIAD and mITD, as a function of the incident sound pressure, intensity, and angle. The Miles model becomes analytically solvable under the assumptions of continuous sinusoidal input and symmetric model parameters, in addition to being numerically solvable without requiring the assumptions of symmetry or continuity. The model allowed Miles *et al.* to demonstrate that *O. ochracea*’s impressive sound localization abilities are due to the pre-processing performed by their structurally coupled tympana, which mechanically amplify the ITD and IAD experienced by the fly.

In addition to providing a physiological explanation for *O. ochracea*’s localization prowess, the Miles model also accurately predicted mITD for all incoming sound angles and mIAD for angles below ±30° in a sample *O. ochracea* population. Both the measured and predicted mITD indicated that *O. ochracea* possesses an mITD comparable to the ITD of an animal closer in size to a rat (Fig. 1B). Later experiments successfully determined that *O. ochracea* has a sound localization precision in the azimuthal plane of 2° (20, 21), a precision comparable to that of humans. This high precision, together with the relative simplicity of the model and the easily reproducible structure of the hearing mechanism used by *O. ochracea* led to a new stream of research in *ochracea*-inspired designs for directional microphones and hearing aids (22–28). Despite its utility, the model contains a number of simplifications that limit its biological accuracy.

The Miles model is a lumped-element model that primarily considers the dynamics of the intertympanal bridge and the front of the tympanal membranes (Fig. 1C), modeling each membrane as a flat plate with a purely one-dimensional amplitude response. The model’s spring and damper coefficients were adjusted until the model response approximated the experimental responses in recently deceased *O. ochracea* specimens measured using laser-vibrometry. Although the model is relatively accurate for mITD in a narrow range of incident sound angles, it displays significant errors in mIAD for incident sound angles larger than approximately ±30° from the midline of the fly, and mITD becomes increasingly inaccurate at angles above approximately ±40°. This inaccuracy across large angles limits the model’s power for explaining binaural hearing in *O. ochracea* and its potential for inspiring new hearing-based engineering.

Previous scanning electron microscopy images of *O. ochracea* tympana had indicated that a certain degree of dynamic curvature and morphological complexity was present (8). It was excluded, however, primarily to avoid increasing model complexity. We hypothesized that inclusion of 3D features could improve model accuracy and extend the effective range of the model, allowing it to predict more accurate tympanal displacements for incoming sound at high angles when compared to experimental data. To identify the sources of inaccuracy in the model, we investigated the detailed morphological structures involved. Using 3D reconstructions of *O. ochracea*’s tympana as a guide, we modified the original model by adding terms that simulate the mechanics of the hearing organ in the lateral plane. We represent these lateral mechanics mathematically via a spatially-dependent asymmetry in the model spring and damper coefficients.

## Materials and Methods

### Synchrotron x-ray imaging of the ormiine tympanal organ

To examine the 3D nature of *Ormia ochracea*’s tympanal morphology, we performed tomographic imaging of preserved *O. ochracea* specimens using the synchrotron x-rays at the Advanced Photon Source at Argonne National Laboratory. Two *O. ochracea* dried specimens were borrowed from the Virginia Tech Insect Collection. The specimens were placed in slender tubes made of polyimide, and the ventral thorax was imaged using beamline 2-BM. Each specimen was imaged using the beamline’s fast 2D phase contrast imaging, giving stacks of images along the *z*-axis at intervals of 1.72 *μm.*

Raw microtomographic images were cropped and downsampled using FIJI (33) and segmented in SlicerMorph, (34) an imaging extension of 3D Slicer. To segment, features of the tympana were highlighted and then rendered in 3D for applicable measurements. The three-dimensional scans are available upon request.

### Previous model

The previous model of binaural hearing in *O. ochracea* includes two components: a mechanical model of the anatomy and a corresponding mathematical model. The mechanical model (15) treats the tympanal structure as a pair of beams pinned at a central pivot, with lumped-mass approximations of the two sides of the hearing organ located at the ends of the beams (Fig. 3A,B). The beams are anchored to the substrate at their distal ends with a pair of symmetric spring-damper elements, and to each other with a third spring-damper element (Fig. 3B). Pressure forces from incident sound waves are applied to the point masses via a forcing function composed of the product of the incident pressure magnitude, the inward-facing unit normal vector, and the tympanal surface area, ***A*** (see the Supplemental Material for numerical values used in this study). A time delay is applied between the left and right sides based on the angle *θ* the incoming sound wave has relative to the midline of the fly, with 0° defined as straight ahead (15).

**Fig. 2.**
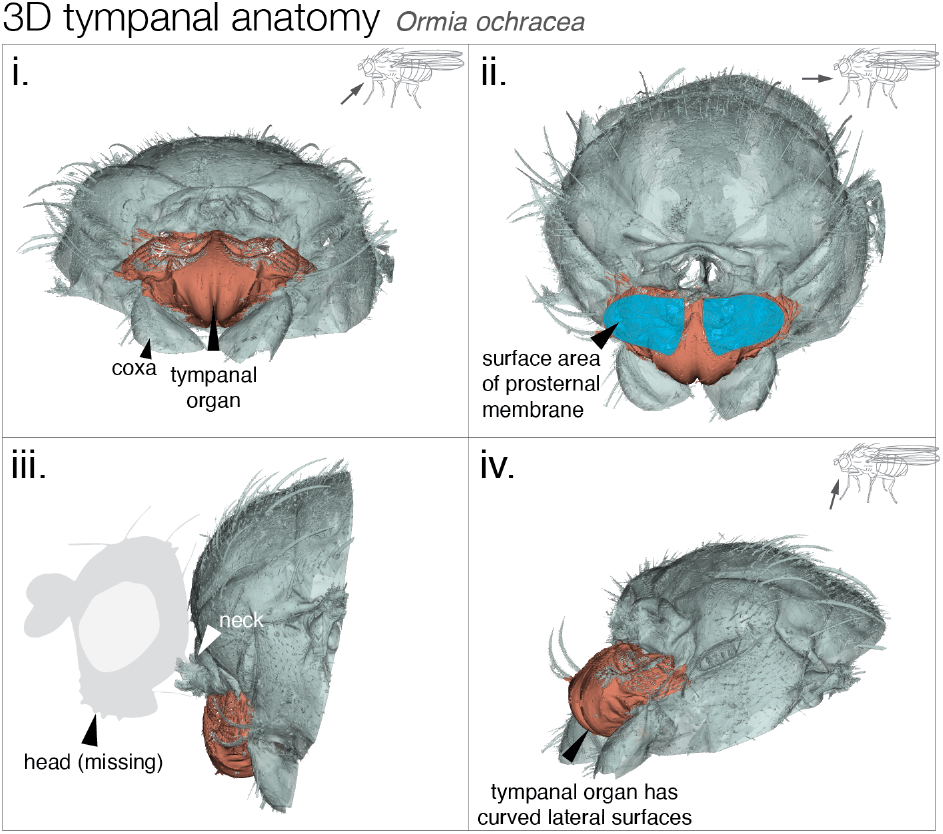
3D rendering of tympanal organs and frontal prothorax of the fly *Ormia ochracea.* Tympanal membranes highlighted in blue (ii.), with supporting structures highlighted in peach. Orientation of image relative to *O. ochracea* body indicated in schematics at top right of images. 3D images made in SlicerMorph software. (34)

**Fig. 3.**
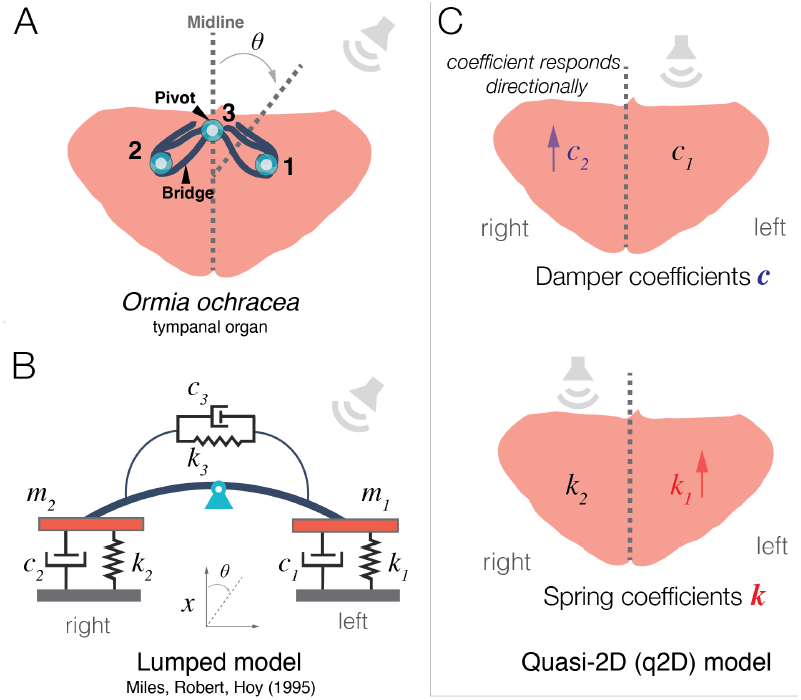
Modeling of binaural hearing in the fly *Ormia ochracea.* **(A)** Schematic of the coupled tympanal membranes of *O. ochracea* (peach-colored, labeled 1 and 2), connected by the cuticular bridge (blue) with sound incident at *θ* degrees. **(B)** The hearing system can be represented as a pair of coupled beams joined and anchored by a set of springs and dampers (adapted from Liu *et al.* (35). **(C)** The q2D model has an asymmetric response: the spring and damper coefficients on the contraleral (opposite) side from the sound source increase as a function of incident sound angle, while the coefficients on the ipsialateral side remain constant.

The mathematical model is a set of coupled ordinary differential equations that are the equations of motion for the mechanical model. It treats the incident acoustic pressure acting on the tympanal membranes as two point forces, ***f***_1_ (***t***) and ***f***_2_ (***t***), acting on the point masses representing the tympanal membranes and associated structures. The dependent variable in the problem is **x**(*t*), which represents the one-dimensional response of each tympanum. The model can be written as:

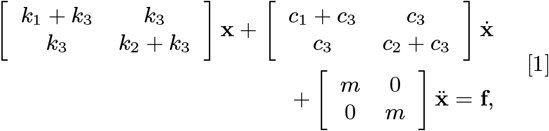

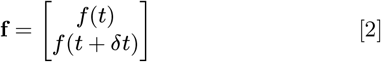

where **x** = (***x***_1_(***t***),***x***_2_(***t***)) is the unknown response vector containing the vertical displacement of the left and rightmost tips of the beams in Figure 3B, which represent the two sides of the intertympanal cuticular bridge, the applied force is **f** = (***f***_1_ (***t***),***f***_2_(***t***)), and 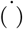 represents differentiation with respect to time, ***t***. The parameters ***k_i_*** and ***c_i_*** are spring stiffness and damper constants, respectively, and the parameter ***m*** is the effective mass of all the moving parts of the auditory system (15).

### Q2D model modifications based on ormiine morphology

In Miles *et al*.’s analysis of their model, the ormiine hearing structure is assumed to be left-right symmetrical, and the spring and damper coefficients on the right and left sides are identical and constant for all incident sound angles, with ***k***_1_ = ***k***_2_ = ***k*** and ***c***_1_ = ***c***_2_ = ***c***, independent of the values of *k_3_* and ***c***_3_.

In order to add a realistic degree of sensitivity to the angle of the incoming sound, we modified the spring and damper parameters to incorporate aspects of the 3D morphology of the fly’s hearing organ. Specifically, we did this by treating the magnitude of ***k*** and ***c*** as functions of the incoming sound angle. The functions were structured such that for an incident sound angle above ±30°, the ***k*** and ***c*** values corresponding to the contralateral tympanum are increased compared to those for the ipsilateral tympanum, mimicking the presence of lateral sides on the tympana, which can both shield the rest of the structure and be more responsive to laterally oriented incoming sounds (Fig. 3C). We provided the following quasi-two-dimensional modification to the Miles model of ormiine hearing:

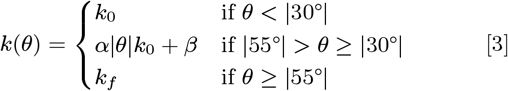

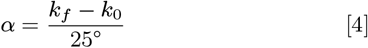

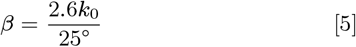

where ***k***_0_ and ***k_f_*** are the minimum and maximum values that the spring stiffness coefficients can take on, respectively. The form of the modified spring coefficient function, two constant segments with a linear ramp between |30|° and |55|° (Fig. 3C, Fig. 4A), was informed by the lateralization behavior observed in *O. ochracea* (20) and the analysis of an *O. ochracea*-inspired sensor (35). These works indicated the presence of two separate behavioral regimes, a localization regime from 0° to ≤ |30|° and a lateralization regime at higher angles. This choice is further supported by the accuracy of the fit to experimental data for sound incident at ≥ |30|° (Fig. 4B,C), and physically represents a degree of elastic response to incoming sound waves in the lateral direction.

**Fig. 4.**
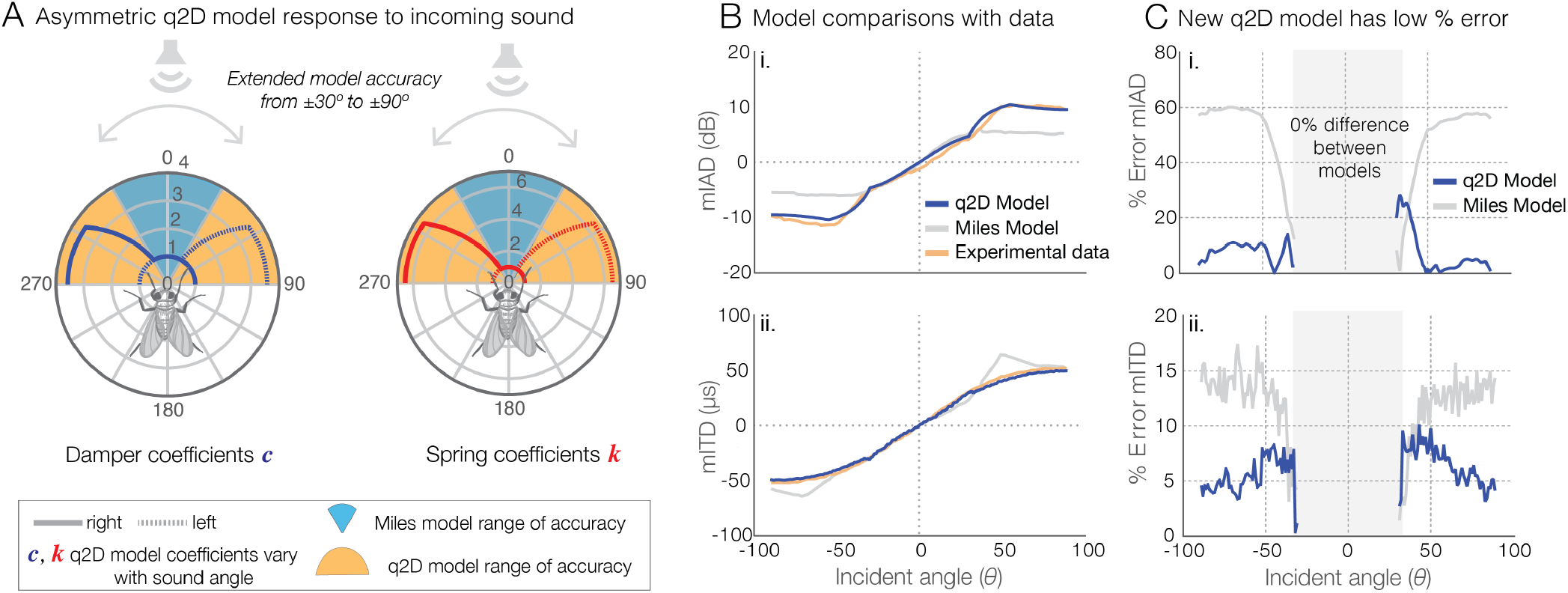
Model modifications and comparison between the Miles model, experimental data, and q2D model. **(A)** The modified quasi-two-dimensional (q2D) model shows improved range of accuracy in its response to incident sound. In the q2D model, the normalized damper (red) and spring (blue) coefficients are functions of the incoming sound angle. The improved q2D model responds accurately within ±90°, compared to ±30° for the Miles model. Experimental and model results **(B)** and error **(C)** in mITD and mIAD for the standard one-dimensional (Miles model) and q2D models. In **(B)**, mITD and mIAD were calculated from the q2D and Miles models as a function of incident sound angle for a frequency of 6000 Hz, and compared with laser-vibrometry measurements from recently deceased *O. ochracea* specimens (15). The significant divergence from behavioral data present in the Miles model outside ±30°, particularly for mIAD, is rectified in the modified q2D model. In **(C)**, the gray box indicates errors below ±30°, which are not considered because the q2D and Miles models are identical for these ranges. The errors for the q2D model peak close to ±30°, then decrease as the incident sound angle is increased.

The constants in equations 3–5 were chosen to provide the best fit to the available behavioral data (15): mITD and mIAD derived from laser-vibrometry measurements of tympanal membrane vibrations in *O. ochracea* specimens in response to a 6 kHz sound source, as a function of incident sound angle. The coefficients are only modified on the contralateral side and remain constant for the side on which the sound source is located. As the incident sound angle approaches ±90° relative to the fly’s head, the spring coefficient for the contralateral side increases from ***k***_0_, and approaches ***k_f_*** according to Equation 3. For example, for sound incident from 30°, the spring and damper coefficients for the left side, ***k***_1_ and ***c***_1_, would change and ***k***_2_ and ***c***_2_ would remain unchanged. We assume the total tympanal surface area, ***A***, is fixed, and we use previously established values (15) throughout this work (***A*** = 0.288×10^-6^*m*^2^). The increases in the spring and damper coefficients, normalized relative to their nominal values (***k***_0_), are visible in Figure 4A.

MATLAB’s ODE45 function was used to integrate equations 1–2 and a custom peak-finding algorithm was implemented to calculate mITD and mIAD. Further computational details and a link to representative code samples can be found in the Supplemental Material.

## Results

The tympana of *O. ochracea* protrude anteriorly from underneath the cervix (fly’s neck), with distinct lateral faces and sharp curvature (Fig. 2). Figure 2 shows 3D surface renderings of *O. ochracea* tympanal membranes in teal, with the supporting structures highlighted in peach. The organs are far from the simple two-dimensional surfaces most often depicted in the literature (14, 15, 28, 35). These new 3D models motivated our modifications to include aspects of actual morphology. The confirmation of significant lateral-facing portions of the tympana led to the modifications present in the q2D model (equations 3–5), which account for the lateral tymapanal response to acoustic stimuli.

Values of mITD and mIAD, calculated from the q2D and Miles models, are shown in Figure 4B as a function of incident sound angle, and are compared to experimental measurements in recently sacrificed *O. ochracea* specimens (15). Both models are identical for incident sound angles less than ±30°, so the results are identical within that range (Fig. 4C, gray box). When we included the lateral response through the new *k*(*θ*) and *c*(*θ*) functions, the gap between experimental measurements and model results in both mIAD and mITD narrowed significantly for 6 kHz signal input (Fig. 4B,C), with the q2D model having average error of approximately 6% and a peak error of approximately 28% in mITD, and an average error of approximately 7% and a peak error of approximately 10% in mIAD. These results additionally confirm that aspects of mechanics in two dimensions are important elements of ormiine hearing.

## Discussion

In this paper, we present the results of 3D X-ray synchrotron imaging of the mechanically-coupled tympana in the parasitoid fly, *Ormia ochracea,* and our subsequent modification to the classic mathematical model of hearing in *O. ochracea* inspired by those results. The tympanal organ was confirmed to be highly 3D, with significant lateral-facing membranes, in contrast to the commonly simplified representation of the membranes as flat, front-facing plates.

Detailed knowledge of the hearing organ’s morphology allowed us to update the classic 1995 one-dimensional mathematical model into a quasi-two-dimensional model of ormiine hearing that mimics the tympanal organ response in the lateral direction. Our updated q2D model has significantly improved fidelity to available experimental data (15) compared to the Miles model, both in the mechanical interaural time delay (mITD) and in the mechanical interaural amplitude difference (mIAD) (Fig. 4B,C). When compared to the Miles model, the new q2D model exhibits maximum errors (relative to experimental values) reduced by approximately 50% and 85% respectively. This strongly supports the premise that there are important aspects of the mechanics of *Ormia* hearing aside from the response of the front-facing tympanal membranes, and that the entirety of the hearing organ structures are sensitive to the angle of incoming sound, a feature that was not included in the Miles model.

Prior to our study, the original Miles model was the only existing model of ICE (internally coupled ears)-based hearing in ormiine flies (36). This is one of the first attempts to update the foundational Miles model for hearing in *O. ochracea.* Our model may be further refined by incorporating additional mechanical behaviors of the tympana, such as tympanal deflection in the lateral direction or a representation of the tympanal response in the vertical direction. It could also be improved by simple analytic modifications to expand the model’s capabilities without impacting its tractability, such as using functions that are more flexible than simple linear ramps for the spring and damper coefficients. For example, in our q2D model, the “bump” visible near ±45° in mIAD in Figure 4B and the uptick at the same point in mITD may be a result of the values for either the springs, dampers, or the ratio between the two, being slightly too high at that point. It is also important to note that this work and the Miles model both rely on tuning the coefficients so that the model outputs better match the experimental response to sinusoidal input (2 kHz for the original 1995 work and 6 kHz for the work here). Although the model’s performance was not observed to degrade at other frequencies that we checked, the degree of improvement (relative to the 6 kHz experimental data) was far less significant for other frequencies. The model’s reduced performance at frequencies other than those tuned specifically for crickets could potentially be resolved by introducing other morphological features in the form of frequency-dependent functions, in a similar way as we have introduced spatially-dependent functions here.

Our model demonstrates that the mechanics of hearing in *O. ochracea* are dependent on the complex tympanal morphology present in the animal, especially with respect to mIAD, and indicates that this morphology serves a specific angle-dependent role in responding to incoming sound waves. The inclusion of angle-dependent behavior in the spring and damper coefficients provides a more accurate understanding of how the insect receives sound. Previous work has demonstrated that *O. ochracea* engages in different behaviors depending on the relative angle of incoming sound (15, 20, 35, 37), with two distinct response patterns. In the first, from 0° to ±30°, the fly makes relatively narrow adjustments to localize the origin of the sound (localization). In the other, at angles exceeding approximately ±30°, the fly makes significantly larger adjustments, more akin to determining the side from which the sound originates (lateralization). Our results show that this difference in response is not strictly a result of behavioral differences, but is paired with a difference in physiological responses to incoming sound.

Furthermore, there is growing evidence that some *O. ochracea* are involved in an evolutionary arms race with their host species (38, 39), and that they are capable of differentiating between different cricket host species based on their acoustic signalling, exhibiting preference towards local populations (40). Consequently, the mechanical parameters for the model may depend heavily not only on the geographic origin of *O. ochracea* samples, but also when collection occurred. The degree of tuning to host-searching behavior, as opposed to predator-avoidance behavior, also remains unaddressed experimentally, despite the startle responses when in flight and subjected to sound consistent with bat sonar frequencies (9). *O. ochracea* also exhibits a sorting behavior (being able to rapidly categorize sounds as belonging to a predator or not) in response to predator-consistent sound sources, as opposed to host or neutral sound sources (9). *O. ochracea* is also only one of many *Ormia* species, which parasitize a diverse range hosts, and display different behavioral responses to the acoustic signalling of their hosts (7). Only *O. ochracea* has been examined in sufficient detail to develop a mechanical model with accurate parameters; consequently, it may be worth investigating the mechanics of other ormiine species (7, 41), and developing mechanical models similar to the q2D model presented here. It may also be worth revisiting the hearing organs in *Emblemasoma,* another group of parasitoid flies, which represent a case of convergent evolution in a distantly related family, *Sarcophagidae* (42, 43).

*O. ochracea*’s hearing system has repeatedly served as a source of inspiration for bio-inspired designs for directional microphones and hearing aids (22–28, 35). Including the angle-dependent behavior of the expanded q2D model in future *Ormia*-inspired device designs may also provide significant avenues for improvement in device performance, or may expand the functionality of devices like acoustic sensors through miniaturization and tunable frequency sensitivities. Currently, work is being undertaken to explore the inclusion of lateral faces on a directional microphone to further study the role that these elements play and to attempt to develop a novel practical application. However, there are numerous avenues for exploration remaining, both experimental and theoretical. These include the development of improved bio-inspired technology by incorporating higher-dimensional features and parameter variations in the mechanical system, studying the behavior of the model at frequencies commensurate with bat sonar, and investigating the role that mechanical differences play in *O. ochracea’s* hearing when addressing acoustic preferences. Finally, our expanded q2D model is the first mathematical model of hearing in an binaural fly that is accurate for all measured incident sound angles. It demonstrates the importance of incorporating higher-dimensional model elements consistent with observed physiology, furthering our understanding of binaural and insect hearing.

## ACKNOWLEDGMENTS

The authors thank the Virginia Tech Insect Collection for lending the *Ormia ochracea* samples for imaging, and Pavel Shevchenko for assistance in imaging at 2-BM at Argonne National Laboratory. This material is based upon work supported by the National Science Foundation under Grant number 2014181.

## Abbreviations

The following abbreviations are used in this manuscript:

ITD: Interaural Time Delay
IAD: Interaural Amplitude Difference (sometimes called the Interaural Intensit
mlTD: Mechanical Interaural Time Delay
mlAD: Mechanical Interaural Amplitude Difference

## Notes

The authors declare no conflict of interest. The funders had no role in the design of the study; in the collection, analyses, or interpretation of data; in the writing of the manuscript, or in the decision to publish the results.

https://github.com/aestaples/Ormia3D

### Competing Interest Statement

The authors have declared no competing interest.

### Summary of Updates

Minor edits to text and figures

