## Supplementary Information for "A biologically accurate model of directional hearing in the parasitoid fly *Ormia ochracea*"

PNAS
www.pnas.org

### Supplementary Information for
A biologically accurate model of directional hearing in the parasitoid fly *Ormia ochracea*

Max R. Mikel-Stites, Mary K. Salcedo, John J. Socha, Paul E. Marek, and Anne E. Staples

Corresponding Author: Anne E. Staples


**This PDF file includes:**

Supplementary Information Text

**Other supplementary materials for this manuscript include the following:**

Sample function .m file used for simulation

**Supplementary Information Text**

The following parameters are those that were used for simulation using ODE45 and MATLAB.

**Input Parameters:**

| | |
|---|---|
| incident sound frequency: | 6000 hz |
| speed of sound in air: | 344 m/s |
| Incident sound amplitude: | 72 dB |

**Physiological Parameters:**

| | |
|---|---|
| mass (m): | 6000 hz |
| distance between membranes (mm): | 0.0012 mm |
| 1995 spring stiffness coefficient ($k_{1,2}$): | 0.576 N/m |
| 1995 spring stiffness coefficient ($k_3$): | 5.18 N/m |
| tympanal surface area (A): | 0.288*10^-6 m^2 |
| 1995 damper coefficient ($c_{1,2}$): | 1.15*10^-5 Ns/m |
| 1995 damper stiffness coefficient ($c_3$): | 2.88*10^-5 Ns/m |
| q2D model maximum stiffness ($k_f$): | 3.5747*$k_1$N/m |
| q2D model maximum damper coefficient ($c_f$): | 6.5*$c_1$Ns/m |

**Mechanical coupling function (separate file).** Sample function to perform mechanical coupling using ODE45 and MATLAB, CouplingFunction.m, available at Staples Lab GitHub page.
